# The Cryo-EM structure of TRPC1 and TRPC5 heterotetramer

**DOI:** 10.1101/2024.11.03.621705

**Authors:** Yixiang Chen, Tong Che, Xinyu Cheng, Xiaoqiang Yang, Ying Fu, Wei Zhang, Sijia Lv, Tingting Yang, Qi Peng, Weiwei Nan, Shuangyan Wan, Yaoguang Hua, Juncheng Li, Xiaoyun Wu, Han Hu, Yuting Zhang, Yinzhen Liu, Mingxing Yang, Shuqi Zeng, Jian Li, Bing Xiong, Jin Zhang

## Abstract

Ion channel heteromers display distinct electrophysiological and ligand-binding properties, allowing precise modulation of cellular functions in specific tissues. Among the TRPC subtypes, the TRPC1/4/5 heteromers are the most abundant isoform in the human brain and represent a primary target for treating anxiety and depression. However, the structural organization of TRPC1/4/5 heteromers remain unclear, limiting the advancement of research on these channels. Here, we report the cryo-EM structure of TRPC1/5 heterotatramer at a resolution of 2.84 Å. It consists of an asymmetric structure with three TRPC5 subunits and one TRPC1 subunit. Further structural and functional analysis revealed that the TRPC1 subunit contributes to an asymmetrical ion conduction pathway, a characteristic that differentiates it from TRPC5 homomers. The unique features of the TRPC1 pore loop and its interface interactions with TRPC5 were shown to play crucial roles in modulating the ion conduction properties and gating mechanisms of the heteromeric complex. In addition, we identified key residues within the gating regions that influence channel activation and ion permeability, elucidating the structural basis for the functional divergence between the TRPC1/C5 heterotetramer and TRPC5 homotetramer. Notably, TRPC5 residues were found to dominate gating mechanisms, underscoring the complex interplay between subunit composition and functional outcomes in TRPC channels. Our findings provide novel insights into the structural and functional dynamics of TRPC1/C5 heteromers, paving the way for targeted therapeutic strategies in TRPC5-related disorders.

## Introduction

The transient receptor potential (TRP) channels, a class of non-selective cation channels located in the plasma membrane, play a crucial role as cellular sensors for pain and temperature in humans (1). Among these, the Transient Receptor Potential Canonical (TRPC) subfamily has garnered significant research interest. This subfamily comprises seven distinct members categorized into two groups based on sequence homology: TRPC1/4/5 and TRPC3/6/7 (2, 3). TRPC channels are widely distributed throughout the human body, with TRPC1/4/5 exhibiting high expression levels in key regions of the brain, such as the hippocampus and amygdala (4–7). Research has demonstrated the existence of TRPC channels in the mammalian brain as four distinct heteromeric complexes: TRPC1/4, TRPC1/5, TRPC1/4/5, and TRPC4/5, with TRPC1/4 and TRPC1/5 being the predominant forms (8). These channels are integral to a spectrum of physiological processes, including neuronal growth, synapse formation, neurotransmitter release, synaptic transmission, and synaptic plasticity (9–11).

Gene knockout studies of TRPC1/4/5 have elucidated their significant roles in the central nervous system. Deficiencies in TRPC4 or TRPC5 in mice correlate with reduced anxiety- and depression-like behaviors, while TRPC4 knockout (KO) rats demonstrate a marked reduction in social exploration behaviors (11–13). In double knockout (DKO) studies, mortality rates following pilocarpine-induced seizures were significantly decreased in TRPC1/4 DKO and TRPC5 KO mice, in contrast to TRPC1 KO mice (14, 15). Furthermore, TRPC1/4/5 triple knockout (TKO) mice exhibited reduced opioid-induced side effects, such as analgesic tolerance and hyperalgesia in response to morphine (16). These findings underscore the essential roles of TRPC1/C4/C5 homomers and heteromers in neuronal function, with individual deficiencies impairing heteromer activity. Recent studies have also identified a TRPC4 splice variant in humans linked to autism, while TRPC5 deficiency has been associated with anxiety, obesity, and autism (17, 18). Collectively, these results indicate that distinct TRPC1/C4/C5 homomers and heteromers play critical and diverse roles in regulating physiological functions.

Early electrophysiological studies utilizing patch-clamp techniques have characterized the ion currents of TRPC4 and TRPC5 homomers, as well as TRPC1/4 and TRPC1/5 heteromers. These studies revealed that TRPC5 and TRPC4 homomers conduct larger inward sodium and calcium currents, producing smaller outward currents at +40 mV. In contrast, the current-voltage profile of TRPC1/4 and TRPC1/5 heteromers resembles that of NMDA receptor channels, displaying a slow inward rectification at depolarized potentials and rapid outward rectification above 0 mV (6). These heteromers exhibit unique biophysical characteristics and demonstrate strong tissue-specific expression in the brain, primarily as TRPC1/4 and TRPC1/5 heteromers (8). Despite extensive research demonstrating the importance of TRPC1/4/5 heteromeric channels in neurons, their structural composition and assembly remain poorly understood, necessitating further investigation into their activation mechanisms.

To address these gaps, we employed single-particle cryo-electron microscopy (cryo-EM) to resolve the structure of the TRPC1/5 heteromeric channel in the apo state at a resolution of 2.84 Å. Our findings reveal that the channel forms a heterotetramer consisting of one TRPC1 and three TRPC5 subunits, exhibiting a pseudo-symmetric tetrameric arrangement with an asymmetric ion pore. Notably, we identified that gating amino acids constituting the asymmetric ion pore significantly influence the functional properties of the heteromer. Additionally, we observed a structurally absent, non-conserved amino acid sequence within the TRPC1 pore loop, which is critical for the unique electrophysiological characteristics of the TRPC1/5 heteromer. Moreover, our study demonstrated calcium ion density at the calcium-binding site of the TRPC5 subunit within the TRPC1/5 heteromer, while no calcium binding was detected at the corresponding position in the TRPC1 subunit, highlighting TRPC1’s role as a negative modulator in heteromer formation (19, 20). We also identified several critical amino acids that influence the functionality and assembly of the heteromer through targeted point mutations at the interaction interface between TRPC1 and TRPC5 subunits. Overall, our research provides a structural basis for a deeper understanding of the functional characteristics of the TRPC1/5 heteromer.

## Results

### Biochemistry and electrophysiology of TRPC1/C5 heterotetramer

Ion channel proteins are typically present in low abundance within organisms, and the isolation of their heteromeric forms poses significant challenges. Our initial goal was to analyze the structure of the human TRPC1/5 heteromer; however, the resolution obtained was insufficient to differentiate between the TRPC1 and TRPC5 subunits. Consequently, we shifted our focus to the heterologous expression of the TRPC1/5 protein using both human and mouse sources, ultimately achieving a cryo-electron microscopy structure of the TRPC1/5 heteromer with a resolution of 2.84 Å (Figure 1C and 1D). To ensure stable expression of the TRPC1/5 heteromeric protein, we inserted the full-length gene sequence of human TRPC1 alongside a truncated gene sequence of mouse TRPC5 (featuring a 210-amino-acid C-terminal truncation) into the pEG BacMam vector (21). Given the inherent instability of TRPC1, we included an MBP (maltose-binding protein) fusion tag at the N-terminus and a GFP-2×Strep tag at the C-terminus to enhance its expression stability. The N-terminus of TRPC5 was similarly tagged with an MBP fusion tag and a 3×Flag tag to facilitate the purification of the TRPC1/5 complex (Figure S1A). We transiently transfected the constructed recombinant plasmid into HEK293T cells and utilized the whole-cell patch-clamp technique to verify that our plasmid construction did not compromise the channel functionality of the TRPC1/5 heteromer or the TRPC5 homomer. As shown in Figure 1A, the ion currents of the heteromeric and homomeric channels expressed by our recombinant plasmid were consistent with those of the wild-type TRPC1/5 and TRPC5 channels.

**Figure 1.**
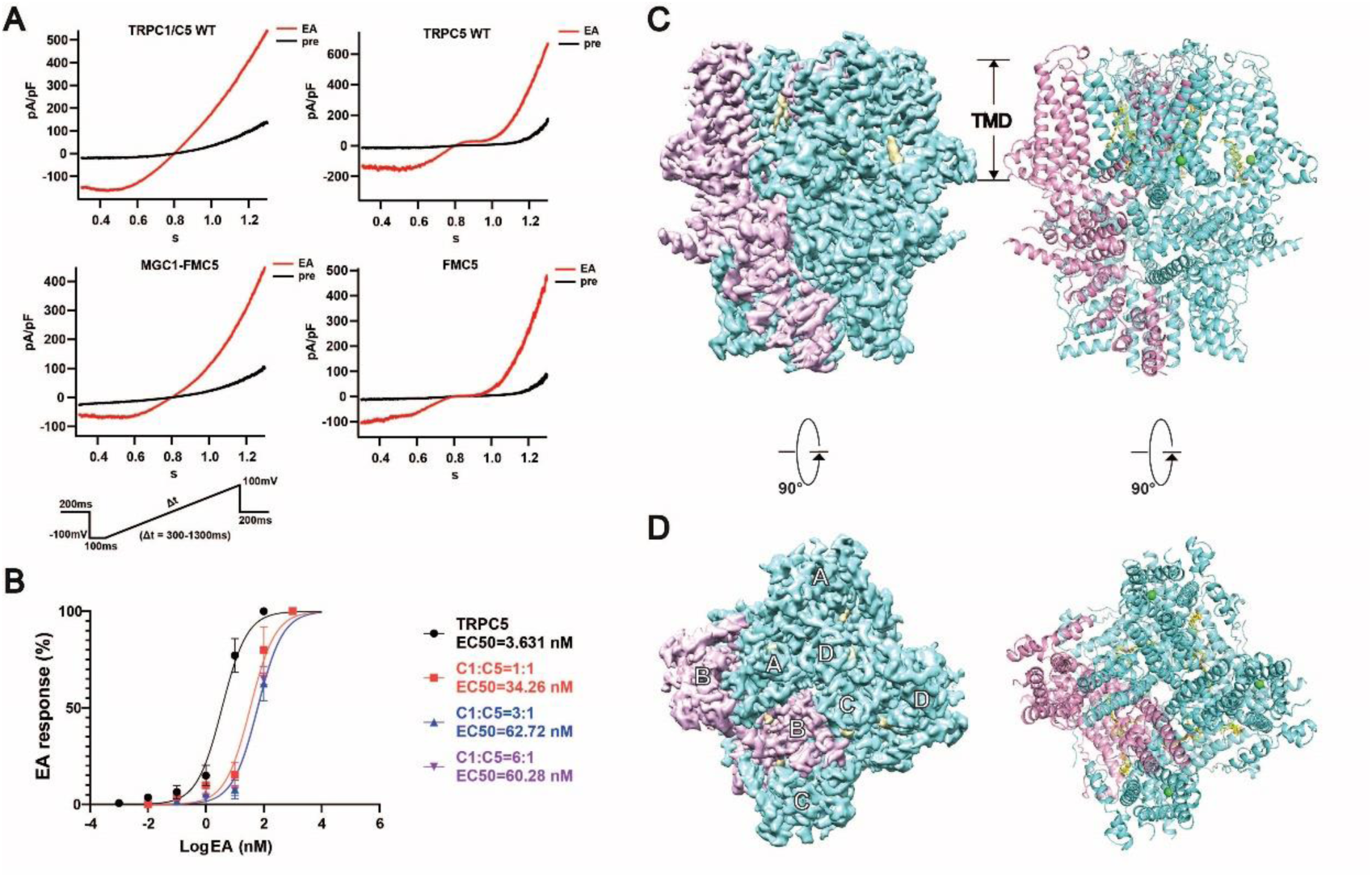
Stoichiometric composition and overall structure of TRPC1/5 heteromer. **(A)** Representative whole-cells currents in HEK293T cells transfected with either full length hTRPC5 (TRPC5 wild type (WT)) or truncated mTRPC5 (FMC5) constructs alone or co-transfected with recombinant plasmids hTRPC1 (MGC1) and mTRPC5 (FMC5) or wild type plasmids full-length hTRPC1 (WT) and full-length hTRPC5 (WT). Currents were recorded before (pre) (black) and after application of 100 nM Englerin A (EA) (red). **(B)** Dose response of EA with EC50 resulting from Hill fits (lines) listed. Data are represented as mean ± SEM (n = 5). **(C and D)** Overall structure of TRPC1/5 heterotetramer. Side and top views of the cryo-EM density map of TRPC1/5 at 2.84 Å overall resolution. TRPC1 subunit represented in pink and TRPC5 subunit represented in aquamarine.

Bernd Fakler and colleagues have previously investigated the composition ratio of TRPC1/C4/C5 heteromer in rat brains, determining that the stoichiometry of TRPC1/5 is approximately 1:3 (8). To explore whether the stoichiometry of the TRPC1/5 heteromer in vitro differed from in vivo expression patterns, we conducted whole-cell patch-clamp recordings to assess the activation of TRPC1/5 heteromer by the TRPC1/C4/C5 non-selective agonist Englerin A at varying transfection ratios. As illustrated in Figure 1B, at a plasmid ratio of 1:1 (TRPC1/TRPC5 w/w), the EC50 was determined to be 34.26 nM, which is closer to that of the TRPC5 homomer (EC50 = 3.631 nM) than to the 3:1 ratio (TRPC1/TRPC5 w/w), where the EC50 was 62.72 nM. This indicates that at the 1:1 ratio, both homomer and heteromer are present; however, the number of homomers diminishes at the 3:1 ratio. When the transfection ratio was further increased to TRPC1/TRPC5 = 6:1 (w/w), the EC50 for heteromer channels activated by Englerin A (EC50 = 60.28 nM) remained nearly unchanged from the 3:1 ratio, suggesting that even with an increased proportion of TRPC1, the activation effect of Englerin A does not enhance (Figure 1B). This observation indicates that at the 3:1 ratio, the majority of channels formed within the cells are indeed TRPC1/5 heteromer, confirming that the stoichiometry of TRPC1/5 is consistent both in vivo and in vitro. This finding provides a solid foundation for subsequent structural analyses; thus, we employed a transfection ratio of 3:1 (TRPC1/TRPC5, w/w) in our subsequent whole-cell patch-clamp experiments.

### Overall structure of TRPC1/5 heterotetramer

We collected over 4,000 cryo-electron microscopy images, and through meticulous data processing and model building, we successfully generated a map of the TRPC1/5 heterotetramer at an overall resolution of 2.84 Å using single-particle cryo-EM (Figure S8). TRPC1 exhibits significant homology with TRPC4 and TRPC5, with amino acid sequence identities ranging from approximately 46% to 48% (https://blast.ncbi.nlm.nih.gov/Blast.cgi). Despite this substantial homology, TRPC1 and TRPC5 can be differentiated based on the electron density of their transmembrane helical side chains (Figure S4). The overall architecture of the TRPC1/5 complex is a heterotetramer, comprising three TRPC5 subunits and one TRPC1 subunit (Figure 1C and 1D). Figures 1C and 1D illustrate the front and top views of the TRPC1/5 heterotetramer, respectively. The TRPC1/5 heterotetramer exhibits structural similarities to the TRPC5 homotetramer, wherein TRPC1 replaces one of the TRPC5 subunits, resulting in a pseudo-symmetric arrangement. Like the TRPC5 homotetramer, the TRPC1/5 heterotetramer consists of six transmembrane helices, with intracellular domains located at both the N-terminus (including the ARD, HLH, and pre-helix domains) and the C-terminus (encompassing the TRP domain and coiled-coil domain). Our findings provide the first structural characterization of the TRPC1 subunit (Figure S1C and S1D). Although the TRPC1 subunit shares structural similarities with the TRPC5 subunit, notable differences are present, particularly in the lengths and tilt angles of certain helices. The S1-S4 helices of TRPC1 are observed to be tilted toward the lower left (Figure S5). The most pronounced distinction is the absence of a segment in the pore loop of TRPC1. Sequence alignment reveals that this segment is not conserved among TRPC1, TRPC4, and TRPC5. Furthermore, the selectivity filter of TRPC1 is not conserved and consists of six amino acids (SLAHVA, residues 600-605), which includes two additional residues, Val and Ala, compared to TRPC4 (GLIN, residues 577-580) and TRPC5 (GLLN, residues 581-584) (Figure 2A, S2). This structural variation contributes to an asymmetric ion conduction pore, which we hypothesize may be a primary factor underlying the outward rectification observed in the TRPC1/5 heteromer.

**Figure 2.**
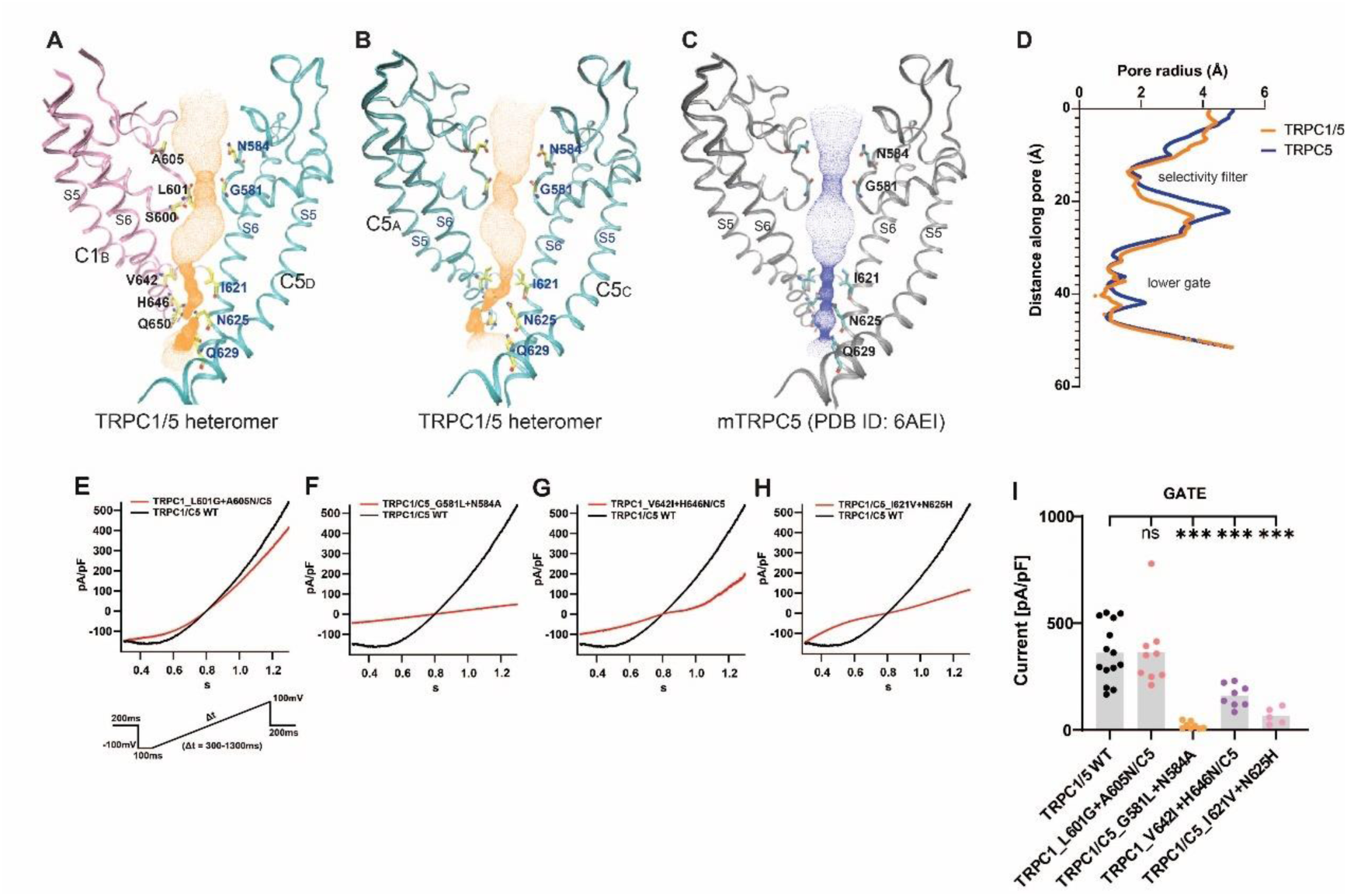
The asymmetric ion-conducting pore of TRPC1/5 heteromer. **(A)** Side view of TRPC1/5’s pore region with chains B (C1, pink) and C (C5, aquamarine). **(B)** Side view of TRPC1/5’s pore region with chains A and C (C5, aquamarine). **(C)** Side view of TRPC5’s pore region with chains A and C (gray). The ion conduction pathway is shown as dots and mapped using HOLE with key amino acid residues labeled. **(D)** Pore radius along the central axis in the apo state of TRPC1/5 heteromer (in orange) and TRPC5 homomer (in dark blue). **(E to H)** Patch clamp recordings of TRPC1/5 mutants in response to channel activators EA. Normalized and superposed current traces of wild-type TRPC1/C5 (WT) (black) and each mutant (red). **(I)** Densities of currents at +80 mV evoked by 100 nM EA for TRPC1/5 WT and mutant constructs. Current density measured as ratio of peak current amplitude to cell membrane capacitance (pA/pF). *p* values were determined by independent t test in SPSS; \**p*<0.05, \*\**p*<0.01, \*\*\**p*<0.001 vs TRPC1/5 WT, each point represents a cell patch.

### Dominant role of TRPC5 in gating TRPC1/5 heteromeric channel

The ion conduction pore of the TRPC1/5 heteromer consists of the pore helix, pore loop, and S5 and S6 helices. In comparison to the TRPC5 homomer, the ion conduction pore of the TRPC1/5 heteromer exhibits notable asymmetry (Figures 2A-D). This asymmetry primarily results from the presence of distinctive gating residues, with both the upper and lower gates of the heteromer being narrower than those of the TRPC5 homomer (Figures 2A-D, S3A-C). Structural analysis identifies key amino acids contributing to the upper gate, specifically L601 and A605 from the TRPC1 subunit, in conjunction with G581 and N584 from the TRPC5 subunit (Figure 2A).

To evaluate the impact of the upper gate on channel functionality, we conducted reciprocal mutations of TRPC1’s L601 and A605 to the corresponding residues G581 and N584 in TRPC5, followed by co-expression assays. The results indicated that the TRPC1^A605N+L601G^ mutation had a negligible effect on the current characteristics of the heteromeric channel. In contrast, the double mutation TRPC5^G581L+N584A^ rendered the heteromeric channel inactive (Figure 2E and 2F). These findings suggest that the TRPC5 subunit plays a dominant role in the gating of the TRPC1/5 heteromeric channel, with G581 and N584 identified as critical residues controlling the upper gate. Previous studies have established the importance of G581 in mediating calcium ion permeability for TRPC4 and TRPC5 (21, 22), a conclusion that is corroborated by our results.

The H646 residue of TRPC1 and the N625 residue of TRPC5 delineate the narrowest region of the lower gate (Figure 2A, S3A). Mutations of TRPC1’s V642 and H646 to the corresponding isoleucine and asparagine residues of TRPC5 resulted in a shift in the current profile of the heteromeric channel from outward rectification to bi-directional rectification, akin to that observed in the TRPC5 homotetramer (Figure 2G). Conversely, mutations of I621 and N625 in TRPC5 to valine and histidine in TRPC1 allowed for activation of the heteromeric channels; however, this was accompanied by a significant decrease in current (P<0.001) and an alteration in current characteristics, with the current transitioning from outward to inward between −100 mV and 0 mV (Figure 2H, 2I).

These findings underscore the critical influence of asymmetric gating residues on the electrophysiological properties of the heteromeric channel, primarily through key amino acids that constitute the lower gate. Additionally, the residues G581 and N584 from TRPC5, which form the upper gate, play a pivotal role in channel activation. Importantly, our observations indicate that mutations of gating residues from the TRPC5 subunit to those present in the TRPC1 subunit significantly impact channel function, further affirming the dominant role of the TRPC5 subunit in the gating mechanisms of the heteromeric channel.

### TRPC1 pore loop: essential for the distinct electrophysiological properties of the TRPC1/5 heteromer

Given the structural similarities between TRPC1 and TRPC5 subunits, what accounts for the marked differences in the electrophysiological properties of TRPC1/5 heteromer compared to TRPC5 homomer? To address this question, we investigated the TRPC1 pore loop and pore helix, which contribute to forming the ion-conducting pore of the heteromer (Figure 3E). Structural alignment revealed a missing segment in the TRPC1 pore loop, which we designated as Loop-T. Sequence alignment further showed that Loop-T is poorly conserved across TRPC1/C4/C5 (Figure 3E and S2). We hypothesized that this segment is critical for TRPC1/5 heteromer activation. To test this, we created two chimeric proteins: TRPC1^Chim-C5^, in which the corresponding TRPC5 segment replaces TRPC1’s Loop-T, and TRPC5^Chim-C1^, where TRPC1’s Loop-T is introduced into TRPC5. When expressed either individually or in co-expression systems, we recorded their currents. Notably, TRPC1^Chim-C5^/C5 currents were nearly identical to those of TRPC5 homomer (Figure 3A), indicating this loop segment as a key determinant of the electrophysiological differences between homomer and heteromer. However, TRPC1^Chim-C5^ alone did not produce any current (Figure 3D), suggesting additional factors, potentially endoplasmic reticulum retention due to its signal peptide, limit TRPC1’s ability to form functional ion channels. Additionally, TRPC1/C5^Chim-C1^ heteromer were non-functional, although TRPC5^Chim-C1^ alone generated bidirectional rectification with a current profile analogous to TRPC5 WT (Figure 3B, 3C and 3F). These findings suggest that the missing flexible region (Loop-T) in TRPC1 is essential for the assembly and activation of TRPC1/5 heteromer but is dispensable for TRPC5 homomer assembly and function.

**Figure 3.**
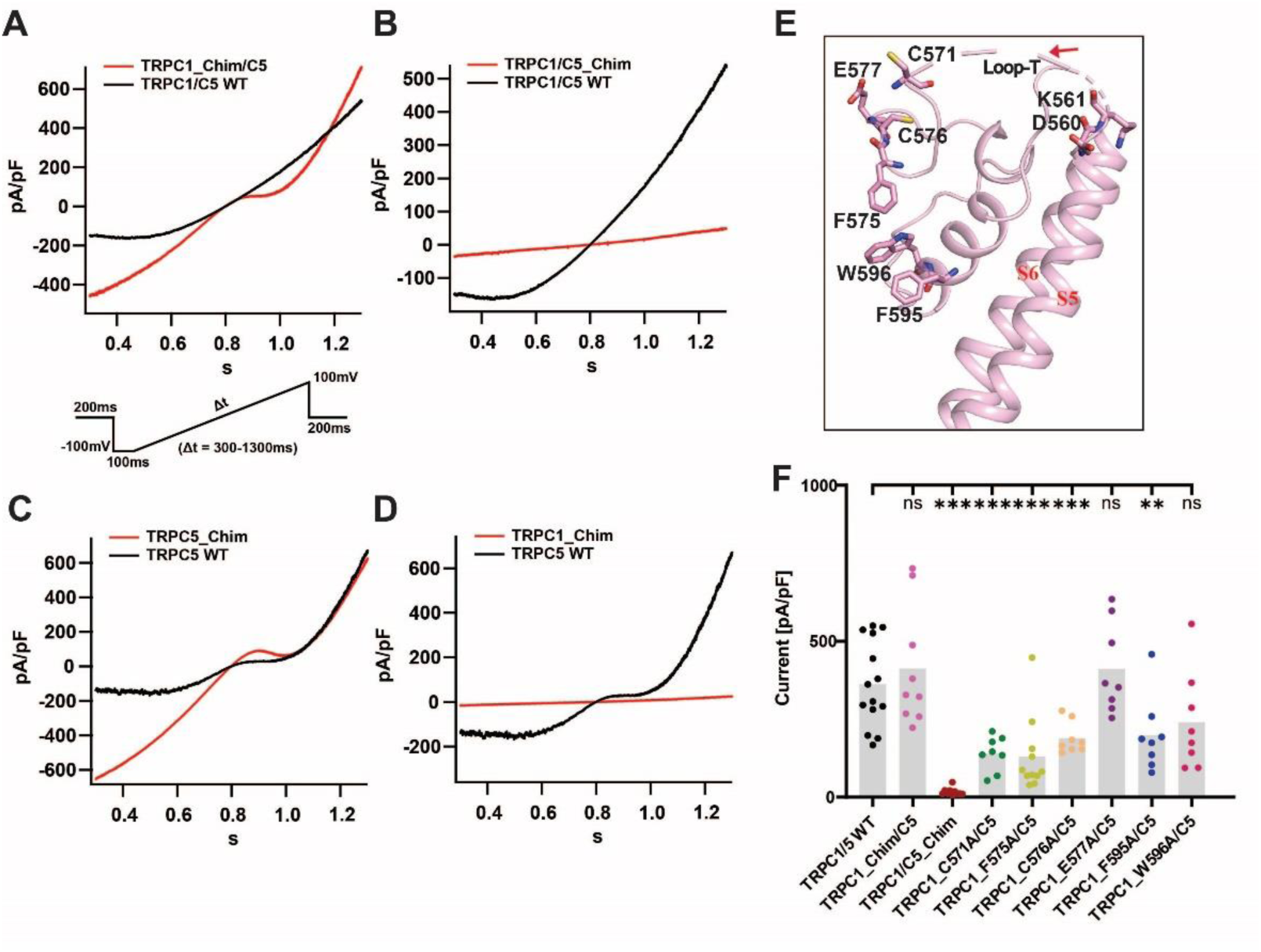
The pore-loop of TRPC1 is essential for TRPC1/5 heteromer. **(A and B)** Patch clamp recordings of TRPC1/5 chimeras (red) and TRPC1/C5 (WT) (black) in response to channel activators EA. TRPC1_Chim/C5 (the corresponding TRPC5 segment replaces TRPC1’s Loop-T (indicated by red arrow) and co-transfected with TRPC5); TRPC1/C5_Chim (the corresponding TRPC5 segment was replaced by TRPC1’s Loop-T (indicated by red arrow) and co-transfected with TRPC1). **(C and D)** Patch clamp recordings of TRPC1 or TRPC5 chimeras (red) and TRPC5 (WT) (black) in response to channel activators EA. **(E)** The pore-helix and pore-loop of TRPC1, the red arrow points to Loop-T. All point mutations of amino acids are displayed in a stick model. (F) Densities of currents at +80 mV evoked by 100 nM EA for TRPC1/5 WT and mutant constructs. Current density measured as ratio of peak current amplitude to cell membrane capacitance (pA/pF). *p* values were determined by independent t test in SPSS; \**p*<0.05, \*\**p*<0.01, \*\*\**p*<0.001 vs TRPC1/5 WT, each point represents a cell patch.

After mutating residues C571, F575, and C576 in the pore loop of TRPC1 to alanine, we observed a significant decrease in the current of the heteromeric channels (P < 0.001) (Figure 3E and 3F). Although the E577A mutation did not affect the current magnitude, it, along with C576A, shifted the current profile closer to that of the TRPC5 homomeric channel (Figure S6C and S6D), indicating that these residues are crucial for the current characteristics of TRPC1/5 heteromer. The conserved LFW motif among all TRPC channels is critical for channel function (7). Notably, double mutations F576A and W577A in TRPC5 resulted in a complete loss of function in the TRPC5 homomer (21, 23). However, mutations F595 and W596 in TRPC1 did not alter the overall activation of the heteromer but shifted its current profile closer to that of the TRPC5 homomer (Figure S6E and S6F). This underscores the dominant role of the TRPC5 subunit in the TRPC1/5 heteromer, highlighting the significance of phenylalanine and tryptophan in the LFW motif in modulating the electrophysiological properties of the heteromer. Our findings suggest that the pore loop and helix of TRPC1 are integral in establishing distinct electrophysiological characteristics compared to TRPC5 homomer, with the absent segment in the TRPC1 pore loop being a critical factor influencing heteromer assembly and electrophysiological changes.

### The interface between TRPC1 and TRPC5

TRPC1 exhibits strong interactions with the adjacent C5 subunits, particularly at the interaction interface between the transmembrane regions. Our investigation focused on the positions labeled B and C in Figure 4A, which highlight magnifications of these critical sites. The S5 and S6 helices of TRPC1 engage with the S4, S5, and S6 helices of the adjacent TRPC5 subunit (Figure 4B), while the S4, S5, and S6 helices of TRPC1 also interface with the S5 and S6 helices of the neighboring C5 subunit on its left (Figure 4C). We introduced mutations of several polar amino acids within this interaction interface to alanine, revealing that the current amplitudes of TRPC1^S619A^/C5^R557A^, TRPC1^K561A^/C5^R390A^, and TRPC1^Q578K+G614R^/C5 significantly decreased relative to the TRPC1/C5 WT (P<0.01, P<0.001) (Figure 4H). Notably, the current profiles of TRPC1^S619A^/C5^R557A^ and TRPC1^K561A^/C5^R390A^ approached those characteristics of C5 homomer (Figure 4D and 4E). These findings suggest that the mutations weakened the interactions between the TRPC1 and TRPC5 subunits, potentially impairing the assembly and activation of the TRPC1/C5 heteromer. Importantly, the polar amino acids involved in these interactions in TRPC1 are not conserved in the TRPC1/C4/C5 complex (Figure S2).

**Figure 4.**
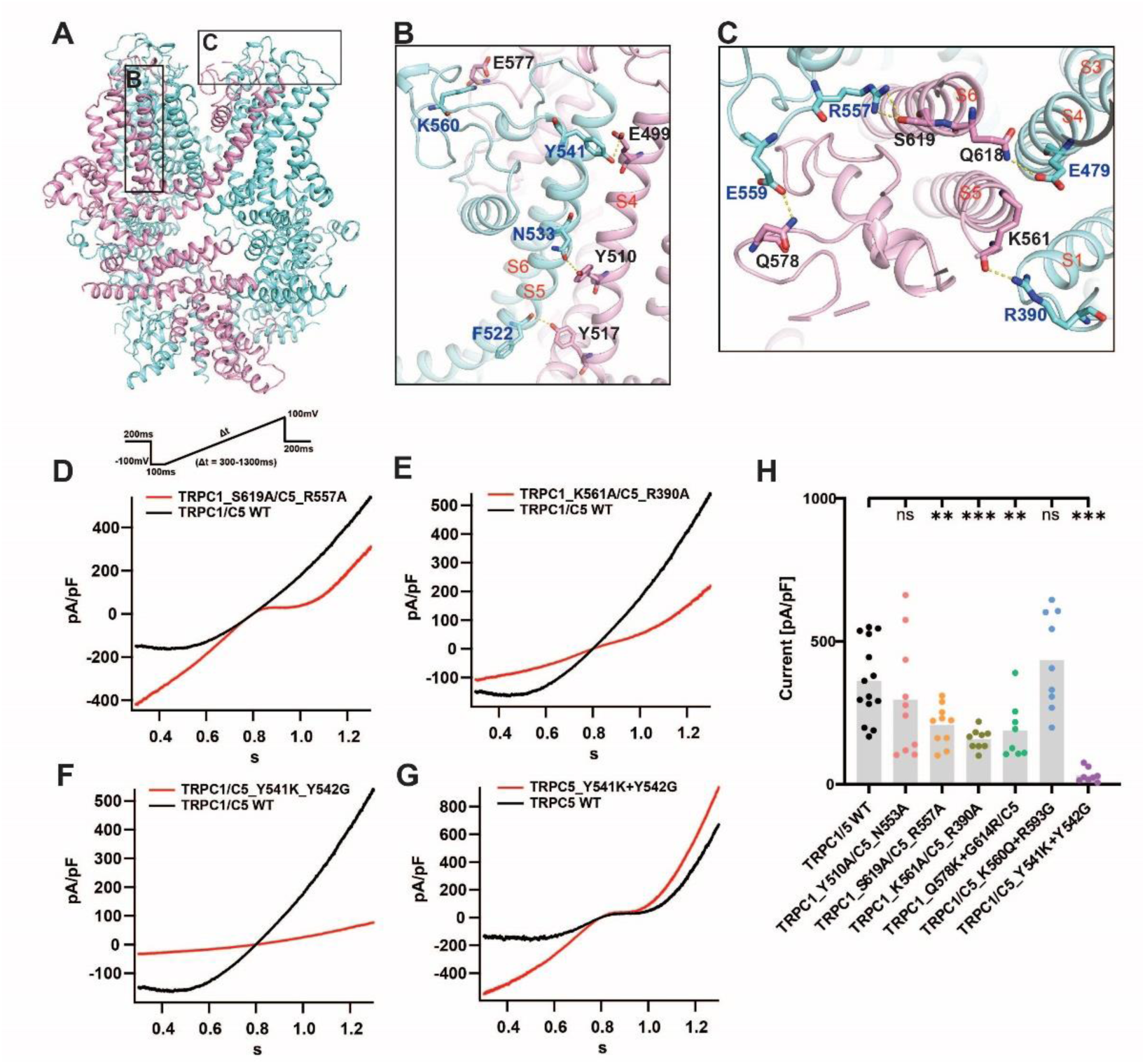
The interaction between TRPC1 and TRPC5 subunits in transmembrane domain. **(A)** Side view of TRPC1/5 heteromer, the two neighbouring TRPC5 subunits and one TRPC1 subunit are coloured aquamarine and pink respectively. **(B)** The amino acids involved in polar interactions between the S5 and pore-loop regions of the TRPC5 subunit and the S4 and pore-loop regions of the TRPC1 subunit. **(C)** The amino acids involved in polar interactions between the S1, S4, and pore-loop regions of the TRPC5 subunit and the S5, S6, and pore-loop regions of the TRPC1 subunit. **(D to G)** Patch clamp recordings of TRPC1/5 mutants in response to channel activators EA. Normalized and superposed current traces of wild-type TRPC1/C5 (WT) (black) and each mutant (red). **(H)** Densities of currents at +80 mV evoked by 100 nM EA for TRPC1/5 WT and mutant constructs. Current density measured as ratio of peak current amplitude to cell membrane capacitance (pA/pF). *p* values were determined by independent t test in SPSS; \**p*<0.05, \*\**p*<0.01, \*\*\**p*<0.001 vs TRPC1/5 WT, each point represents a cell patch.

Furthermore, we identified key amino acid residues at the interaction interface that influence the normal function of heteromeric TRPC channels. Sequence alignment revealed that TRPC1’s K561 and G562 correspond to TRPC5’s Y541 and Y542, a pattern also observed in TRPC4 (Figure S2). Notably, mutating the TRPC5 subunit’s Y541 and Y542 to lysine and glycine in the TRPC1 subunit disrupted the interaction between Y541(TRPC5) and E499(TRPC1). Patch-clamp recordings demonstrated that the heteromeric channel current was barely detectable under these conditions. Conversely, the TRPC5^Y541K+Y542G^ double mutant homomer exhibited normal current consistent with TRPC5 WT, suggesting that the inability of TRPC1 to function as a channel may stem from this disrupted interaction (Figure 4B, 4F, and 4G).

### Ca2+ binding site in TRPC1/5 heterotetramer

The E418, E421, D439, and N436 residues located in the S2 and S3 segments of TRPC5 constitute the calcium ion binding site. Aspartic acid and glutamic acid are predominantly negatively charged, facilitating the attraction of calcium ions to this site (24). Our heteromeric structure reveals a similar binding pattern, with each TRPC5 subunit coordinating a calcium ion at its designated binding site. In contrast, no calcium ion binding was observed at the analogous position in the TRPC1 subunit (Figure 5C and 5D). The four amino acids at the specified positions in TRPC1—D438, R441, N456, and S459—differ from those in TRPC4 and TRPC5 (Figure 5C, 5D, and S2). Notably, R441 is positively charged, while S459 is uncharged, leading to an insufficient negative charge in the binding pocket to attract calcium ions. We introduced mutations in TRPC1, substituting R441 and S459 with Glu (E) and Asp (D), respectively; however, these alterations did not produce significant changes relative to the wild type (Figure 5A). In contrast, mutating E421 and D439 in TRPC5 to the corresponding residues found in TRPC1 preserved the current shape, yet resulted in a significant reduction in current magnitude (P < 0.01) (Figure 5B). This analysis parallels findings from the previously studied TRPC5-riluzole structure, where the hTRPC5 triple mutant (E418Q, E421Q, and D439N, referred to as EED mutants) significantly diminished TRPC5 activation in response to 14 mM extracellular calcium (25). This observation reinforces the notion that TRPC5 plays a predominant role within the heteromeric complex. However, in the case of the TRPC5^E421R+D439S^ mutation, while channel current is not entirely abolished, a marked reduction is evident, suggesting a diminished permeability to calcium ions. This further explains structurally that TRPC1 acts as a negative regulator in the formation of heteromers.

**Figure 5.**
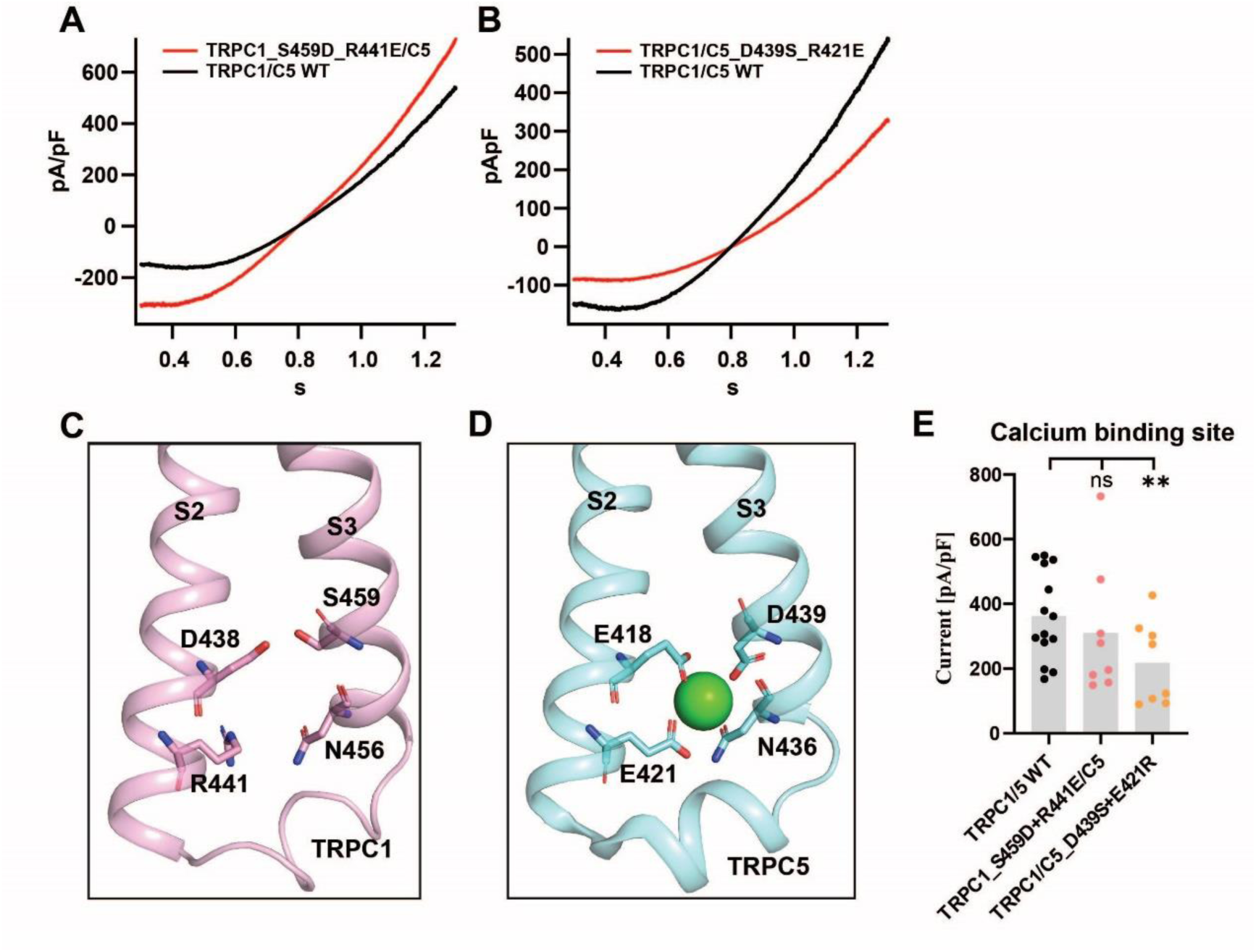
The Ca^2+^ binding site in TRPC1/5 heterotetramer. **(A and B)** Patch clamp recordings of TRPC1^S459D+R441E^/C5 and TRPC1/C5^D439S+R421E^ heteromer (red) and TRPC1/C5 (WT) (black) in response to channel activators EA (100 nM). **(C)** The cartoon representation of S2 and S3 helices of TRPC1. There is no Ca^2+^ binding in this pocket formed by D438, S459, R441, and N456 of TRPC1 subunit. **(D)** The cartoon representation of calcium binding site formed by E418, D439, E421, and N436 of TRPC5 subunit, showing calcium binding. **(E)** Densities of currents at +80 mV evoked by 100 nM EA for TRPC1/5 WT and mutant constructs. Current density measured as ratio of peak current amplitude to cell membrane capacitance (pA/pF). *p* values were determined by independent t test in SPSS; \**p*<0.05, \*\**p*<0.01, \*\*\**p*<0.001 vs TRPC1/5 WT, each point represents a cell patch.

## Discussion

An important hallmark of TRPC channels is their subtype diversity, resulting from various combinations of subunits that form receptor ion channels with distinct electrophysiological profiles, activation mechanisms, compound binding affinities, and spatio-temporal expression patterns. Using cryo-electron microscopy, we elucidated the protein structure of the TRPC1/5 heteromer. Our structural analysis identified a non-conserved amino acid sequence in the pore loop of the TRPC1 subunit, which may influence both the assembly of the TRPC1/5 heteromer and its channel activation. Additionally, we identified several critical amino acids and polar interactions that modulate the functional dynamics of the channel.

Previous studies have suggested that inhibiting or knocking out TRPC5 can alleviate anxiety- and depression-like behaviors in mice (11). However, recent findings indicate that specific knockout of TRPC5 in oxytocin-producing neurons (POMC) leads to obesity and postpartum depression, while TRPC5 overexpression appears to mitigate these symptoms (18). This underscores the dual roles that TRPC5 channels may play, contingent on the specific neural pathways involved. Moreover, TRPC5 knockout partially reduces the expression of TRPC1/5 heteromer, highlighting the complexity of neural signaling.

Given this intricacy, it is imperative to develop compounds that specifically target TRPC subtypes to optimize therapeutic efficacy while minimizing side effects associated with distinct homomeric and heteromeric channels. This necessitates the creation of selective modulators that target TRPC1/5 heteromer while sparing homomeric TRPC5, which is crucial for avoiding off-target effects and maximizing therapeutic efficacy in neuropsychiatric disorders. The structural insights gained from our study of the TRPC1/5 heteromer will pave the way for the development of molecules with heteromer-selective properties.

## STAR ★ Methods

### Method details

#### Plasmid construction

A C-terminal truncated 210 residues amino acid sequence of mouse TRPC5 was cloned into the pEG-BacMam vector. For structural analysis, we designed triple Flag tag with a maltose binding protein (MBP) on its N terminus. A full-length amino acid sequence of human TRPC1 with N-terminal MBP and C-terminal eGFP followed twin Strep-tag®Ⅱ was also cloned into the pEG-BacMam vector. For electrophysiological experiments, the full-length human TRPC5 or C-terminal truncated 210 residues amino acid sequence of mouse TRPC5 with N-terminal mCherry was cloned into pcDNA4.0 vector. All mutants in this study were generated from the recombinant plasmids mentioned above through the design and use of modified primers.

#### Expression and purification of TRPC1/5 heteromer

The expression and purification of TRPC1/5 heteromer were performed as previously described with slight modification (21). Briefly, plasmids FMC5 and MGC1 were respectively transformed into *E. coil* DH10Bac competent cells to obtain bacmids. P4 baculovirus were produced in the Bac-to-Bac Baculovirus Expression System (Invitrogen). HEK293S GnTI^−^ (from the American Type Culture Collection (ATCC)) cells were infected with 10% (v/v) P4 baculovirus with a ratio of TRPC1 to TRPC5 of 2:1 (v/v) at a density of 2.0-3.0*10^6^ cells/ml. Following an incubation period of 18-24 hours, sodium butyrate was administered to enhance protein expression, which was subsequently induced at 30℃ for 48 hours. Then cells were harvested and frozen at −80℃. The cell pellet was re-suspended in buffer A (50 mM Hepes pH 7.5, 150 mM NaCl, and 1 mM DTT, EDTA-free protease inhibitor cocktail (Roche), 10% (v/v) glycerol, 0.5% (w/v) LMNG) (Anatrace), 0.1% (w/v) CHS (Anatrace)) and then solubilized for 3 hours in 4℃. Then the cell lysate was centrifuged for 60 min at 100,000g, and the supernatant was incubated with Strep-Tactin resin (IBA) at 4°C for 2 hours. The resin was washed once with buffer B (50 mM Hepes pH 7.5, 150 mM NaCl, and 1 mM DTT, 10% (v/v) glycerol, 0.1% (w/v) LMNG) (Anatrace), 0.02% (w/v) CHS (Anatrace)), followed by one wash with buffer C (50 mM Hepes pH 7.5, 150 mM NaCl, and 1 mM DTT, 10% (v/v) glycerol, 0.03% (w/v) LMNG) (Anatrace), 0.006% (w/v) CHS (Anatrace), 0.01% (w/v) GDN (Anatrace)). The protein was eluted with buffer C plus 5mM desulfurized biotin (Sigma-Aldrich) and then incubated with anti-Flag M2 affinity resin (Genescript) at 4°C for 1 hours. The resin was washed twice with buffer C and the protein was final eluted with buffer C plus 300 μg/ml Flag peptide (Genescript). The protein was further purified on a Superose 6 (10/300GL) size exclusion column in SEC buffer (25 mM Hepes pH 7.5, 150 mM NaCl, and 1 mM DTT, 0.00675% (w/v) LMNG) (Anatrace), 0.00135% (w/v) CHS (Anatrace), 0.00225% (w/v) GDN (Anatrace)). The peak fraction was collected and concentrated to 8.5 mg/ml for Cryo-EM analysis.

#### Cryo-EM sample preparation and data acquisition

3μL of the samples were applied to Quantifoil grid R1.2/1.3 Au 300 mesh grids (glow discharged at 15mA for 40 seconds with a Glow discharge cleaning system). Grids were blotted with qualitative filter paper in a Vitrobot Mark Ⅳ (Thermo Fisher Scientific) at 4℃ and 100% humidity for 3.5 seconds using a blot force of 2 prior to plunging into liquid ethane and stored in liquid nitrogen until checked.

We used the 300 kV Titan Krios Gi3 microscope (Thermo Fisher Scientific FEI, the Kobilka Cryo-EM Center of the Chinese University of Hong Kong, Shenzhen) to check the grids and collect at 105,000x magnification (pixel size of 0.83Å/pixel). The movie stacks were automatically acquired with the defocus range from −1.0 to −2.0 μm. Micrographs were collected with a total dose of ~54 e−/Å2. SerialEM 3.7 was used for semiautomatic data acquisition. Summary of detailed data collection was shown in Table S1.

#### Cryo-EM data processing

A total of 4,095 movies stacks were imported into cryoSPARC v4.1.1. After motion corrected, electron-dose weighted and CTF estimation, the initial particles were performed by cryoSPARC blob picker. After four rounds of 2D classification, the good particles proceeded to two rounds of Ab-initio reconstruction and heterogeneous refinement. Then we performed an additional round of 2D classification using the best class of particles and selected the template. And the particles were performed by cryoSPARC template picker. After four rounds of 2D classification, the good particles proceeded to two rounds of Ab-initio reconstruction and heterogeneous refinement. We merged the particles of the two best classes to remove duplicates and the final particle sets were re-extracted with original box size and further applied for final nonuniform refinement and local refinement a density map was obtained with overall resolution of 2.84Å (determined by Gold-standard Fourier shell correlation (GSFSC) using the 0.143 criterion).

#### Model building and refinement for cryo-EM structures

The reference models, PDB 6AEI (mTRPC5) and AF-P48995-F1 (predicted structure of hTRPC1) (21, 26, 27), were initially fitted into the EM density map as rigid bodies using Chimera (28). This fit was further optimized using the jiggle fit function in Coot (29), followed by manual adjustments with the real-space refine zone function in Coot to produce an atomic model. Subsequently, the model was refined with the real_space_refine tool in Phenix (30). Validation was performed using MolProbity and Mtriage. PyMOL and Chimera were employed for additional structural analysis and figure generation.

#### Electrophysiology

The full-length or truncated TRPC5, and the full-length TRPC1 constructs was respectively co-transfected in a plasmid ration 1:3 (w/w) into HEK293T cells (from the ATCC). As previously described, a mCherry was inserted into TRPC5 constructs and a GFP was inserted at the C terminus of TRPC1. Therefore, Cells with red and green fluorescence were selected for wholecell patch recordings (HEKA EPC10 amplifier, PATCHMASTER software (https://www.heka.com/downloads/downloads_main.html#down_patchmaster)). A 1-s ramp protocol from −100 to +100 mV was applied at a frequency of 0.2 Hz. Signals were sampled at 50 kHz and filtered at 2.9 kHz. The pipette solution contained 130 mM CsCl, 1 mM MgCl_2_, 5.7 mM CaCl2, 10 mM EGTA, and 10 mM Hepes (calculated free Ca^2+^, 200 nM), and the pH was titrated to 7.2 using CsOH. The standard bath solution contained 140 mM NaCl, 5 mM KCl, 1 mM MgCl_2_, 2 mM CaCl_2_, 10 mM Hepes, and 10 mM D-glucose, and the pH was adjusted to 7.4 with NaOH (1M). The recording chamber (150 μl) was perfused at a rate of ~2 ml/min. All recordings were performed at room temperature. In this experiment, TRPC5 homomer or TRPC1/5 heteromer were activated with the TRPC1/4/5 agonist Englerin A (EA), to assess their electrophysiological properties.

#### Quantification and statistical analysis

Data analysis was performed using GraphPad prism 10.1.2 and IBM SPSS statistics 27. The electrophysiological data were expressed as mean ± SEM, n represents number of independent measurements. Differences between wild type and mutants were evaluated by independent samples T-Test. \**p*<0.05; \*\**p*<0.01; \*\*\**p*<0.001; and ns, *p*>0.05.

## Acknowledgments

Our work was supported by School of Basic Medical Sciences, NanchangUniversity. We would like to thank the Cryo-EM center of The Chinese University of Hong Kong and Shuimu BioSciences Ltd. for the cryo-EM grids screening and data collection. J.Z. was supported by the National Natural Science Foundation of China (grant no. 32271260), the CAS “Light of West China” Program (xbzg-zdsys-202005) and the Jiangxi Province Natural Science Foundation (grant no. 20224ACB206046). J.L. was supported by the Open Project of Key Laboratory of Prevention and treatment of cardiovascular and cerebrovascular diseases, Ministry of Education (No. XN201904), Gannan Medical University (QD201910), Jiangxi key research and development program (20203BBG73063) and Jiangxi “Double Thousand Plan”. T.C. was supported by Jiangxi Provincial Natural Science Foundation (grant no. 20242BAB20119).

## Author contributions

Y.C. performed the experiments (including designing and generating all recombinant DNA plasmids and mutants, optimizing expression and purification of TRPC1/5 complexes, determining the electrophysiological function of TRPC1/5 heteromer), interpreted the data, prepared the figures, and wrote the first draft of the manuscript (ms). J.L. contributed to building the atomic model on the basis of cryo-EM maps and revised the ms; X.C. carried out cryo-EM data processing; H.H., X.W. and Y.Z. performed functional studies. Y.Z. contributed to cryo-EM data acquisition; X.Y., Y.L.and M.W. performed animal studies; W.N. contributed to the expression of Sf9 cells and the harvesting cells; J.Z., B.X. and J.L. initiated the project, conceived the study, designed the experiments, interpreted the data, wrote the ms and supervised for the research. The other authors all coordinated the study.

## Supplementary information

**Figure S1.**
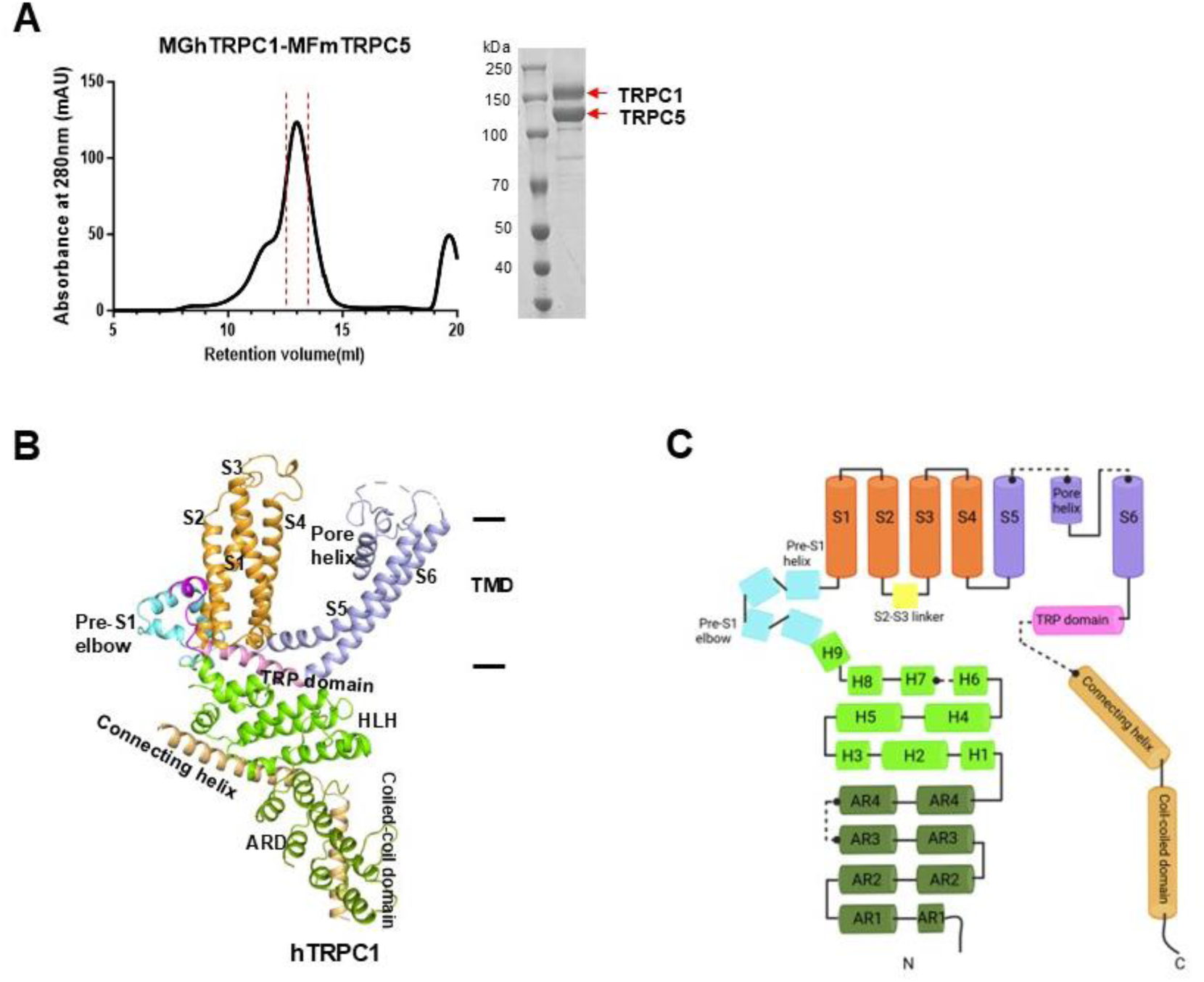
Expression and purification of TRPC1/5 heteromer. **(A)** Size exclusion chromatography of TRPC1/5 heteromer. The peak fractions of TRPC1/5 (between the two red dashed lines) were collected and concentrated for cryo-EM studies. **(B)** Summary of TRPC1/5 heteromer cryo-EM construct design. **(C)** Ribbon diagrams depicting structural details of TRPC1 subunit. **(D)** Linear diagram depicting the major structural domains of the TRPC1 subunit, color-coded to match the ribbon diagram in (C). AR, ankyrin domain; HLH, helix-to-helix.

**Figure S2.**
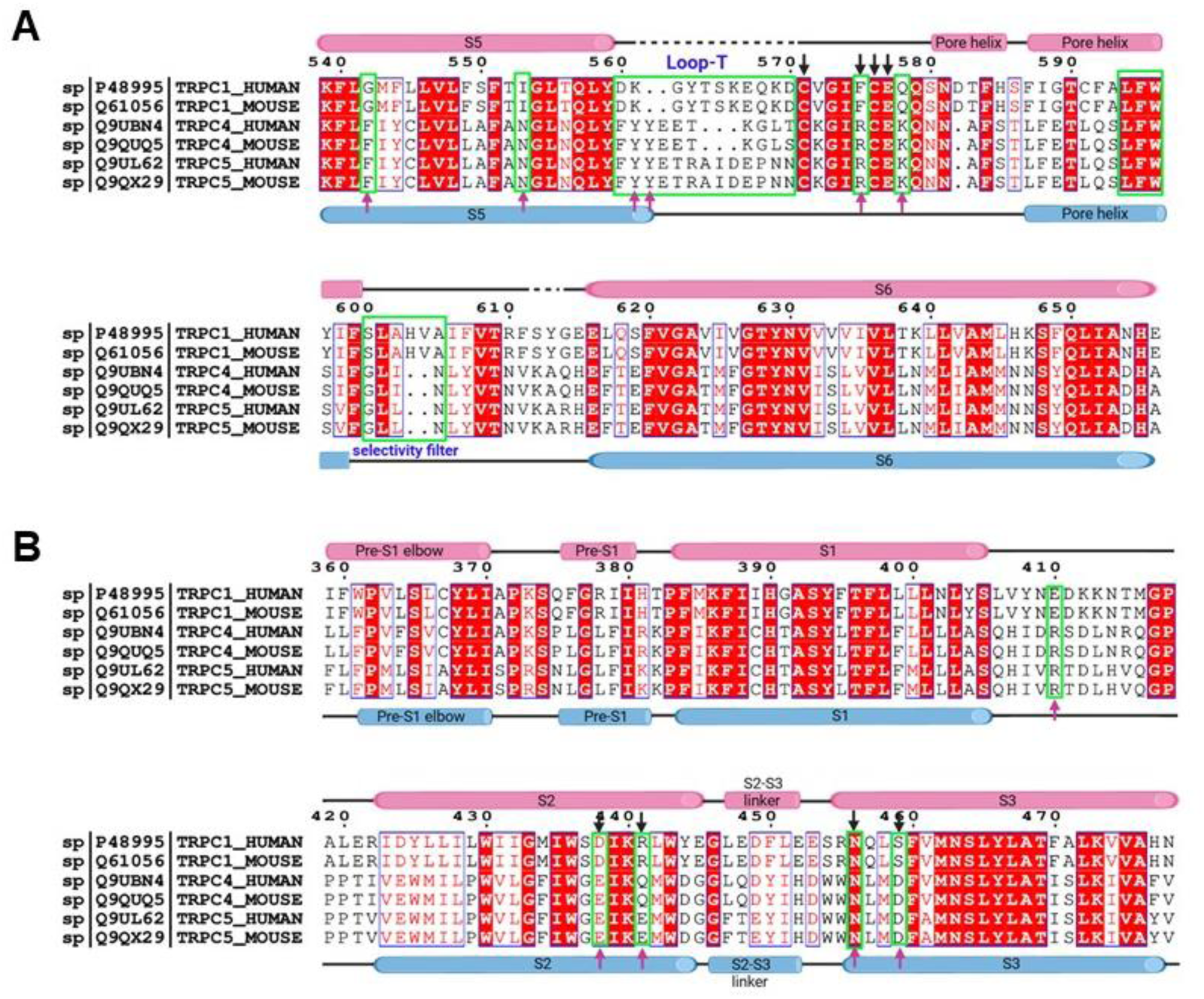
Sequence alignment between human and mouse TRPC1/C4/C5. Secondary structure assignments are based on the structure of apo TRPC1/5, TRPC1 is pink and TRPC5 is blue. The black arrows indicate the residues of TRPC1, and the red arrows indicate the residues of TRPC5. **(A and B) Related to Figure 3, Figure 4, and Figure 5**.

**Figure S3.**
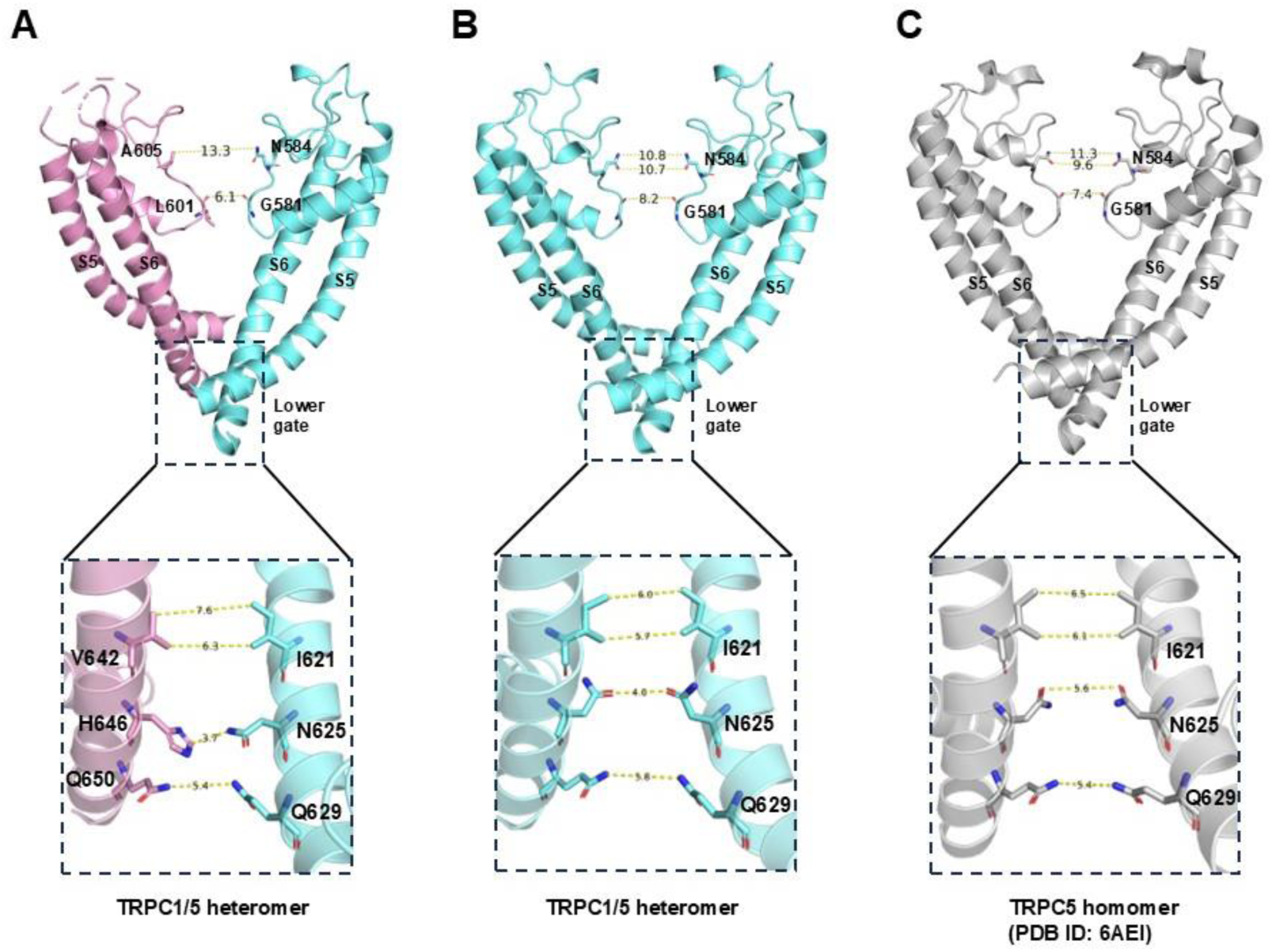
Comparison of the upper gate and lower gate between the TRPC1/5 heteromer and TRPC5 homomer. **(A)** Side view of TRPC1/5’s pore region with chains B (TRPC1 colored by pink) and D (TRPC5 colored by aquamarine). The side chains of L601 (TRPC1) and G584 (TRPC5) form a narrow constriction at the selectivity filter. “VHQ” motif of TRPC1 and “INQ” motif of TRPC5 forms the lower gate, and the narrowest point is between H646 and N625. **(B)** The side view of the ion conduction pore between the two TRPC5 subunits (chains B and D) in the TRPC1/5 heteromer. **(C)** Side view of TRPC5’s pore region with chains A and C (gray).

**Figure S4.**
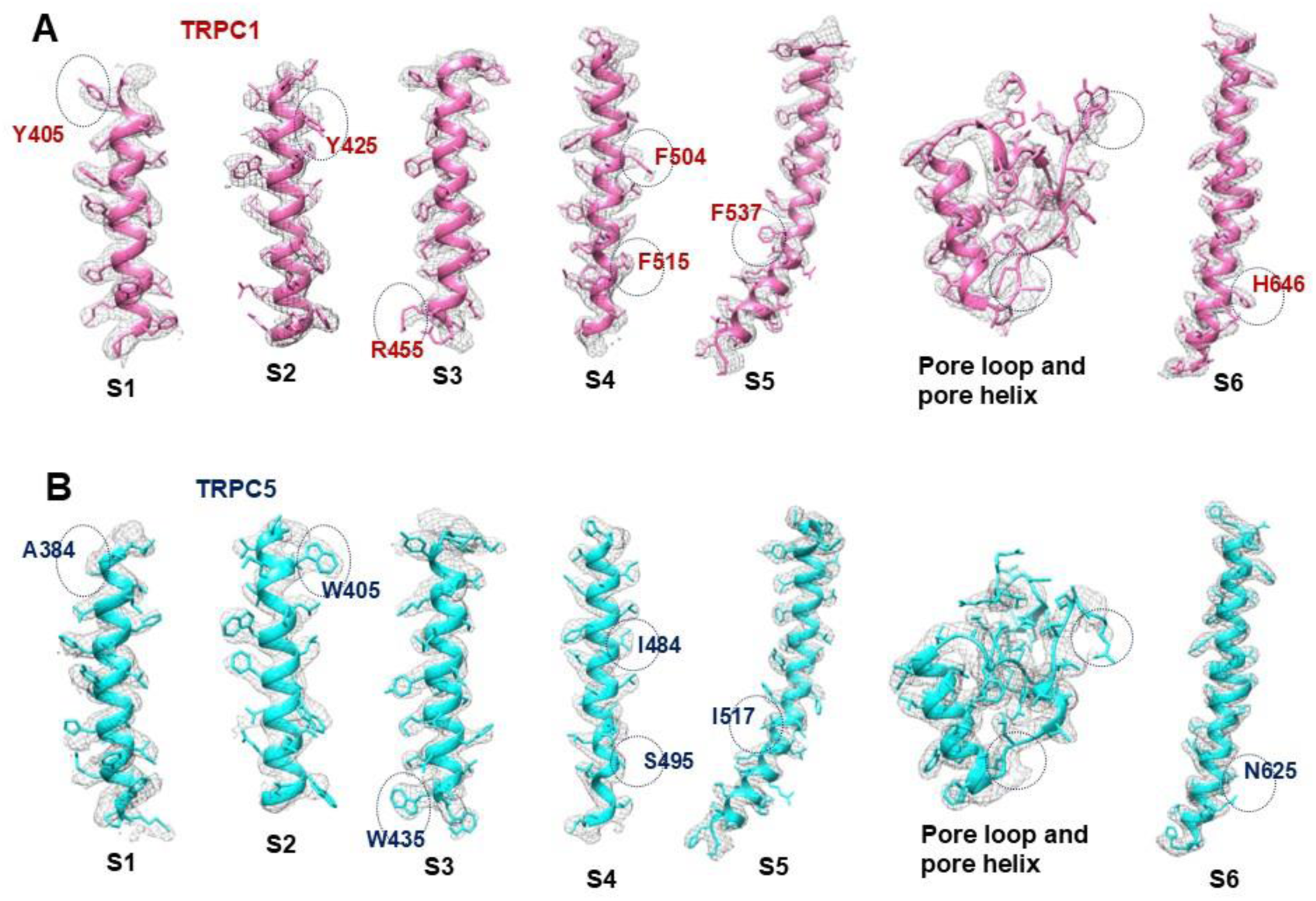
Cryo-EM density for TRPC1/5 heteromer. **(A)** Cryo-EM densities for each segment of the transmembrane helices of TRPC1. **(B)** Cryo-EM densities for each segment of the transmembrane helices of TRPC5. The black dashed circle indicates the different residues or regions between TRPC1 and TRPC5 in cryo-EM density maps.

**Figure S5.**
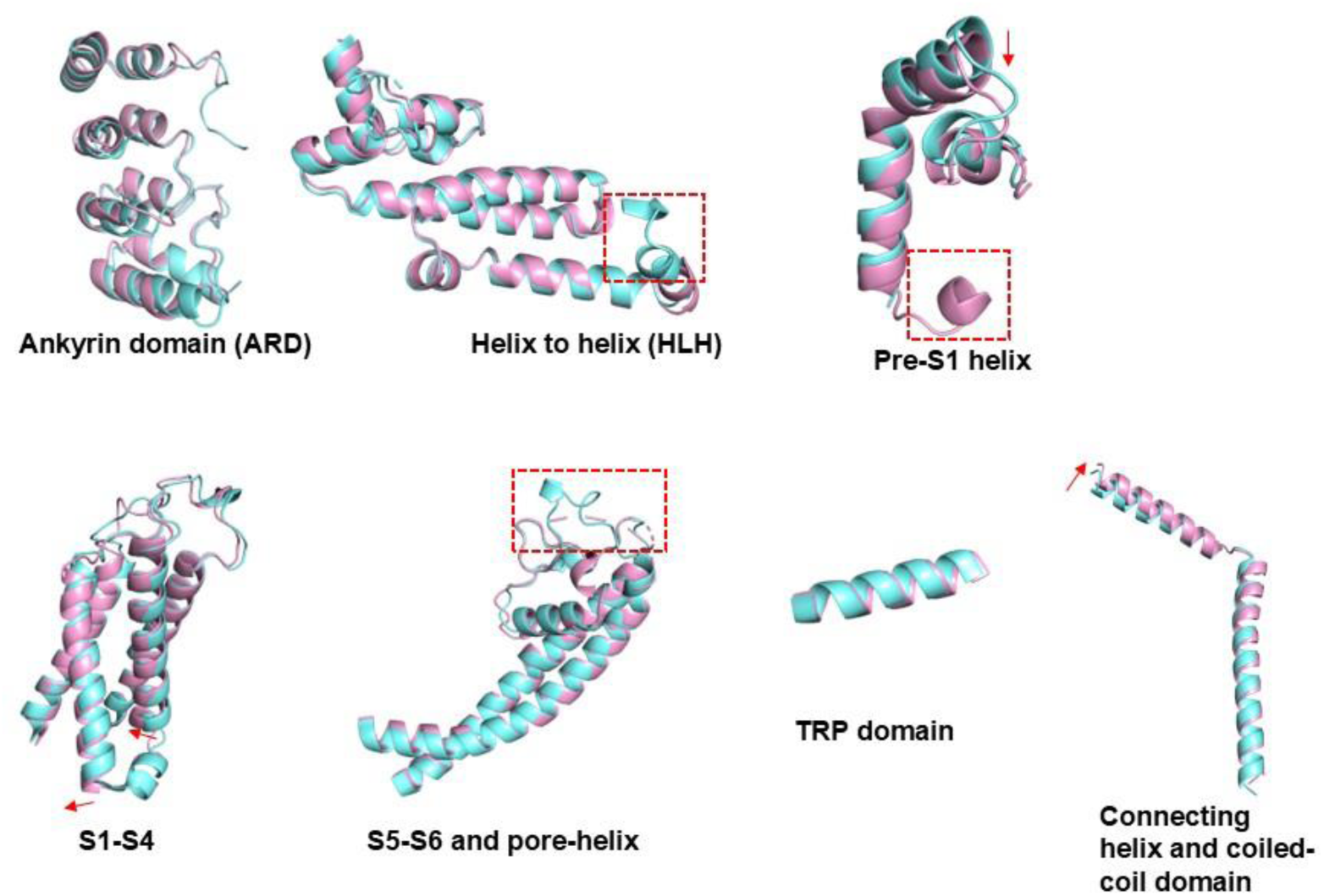
Structural comparison between the TRPC1 (pink) and the TRPC5 (aquamarine). The red arrows indicate the direction of the helix rotation. The red dashed box highlights structural differences between TRPC1 and TRPC5.

**Figure S6.**
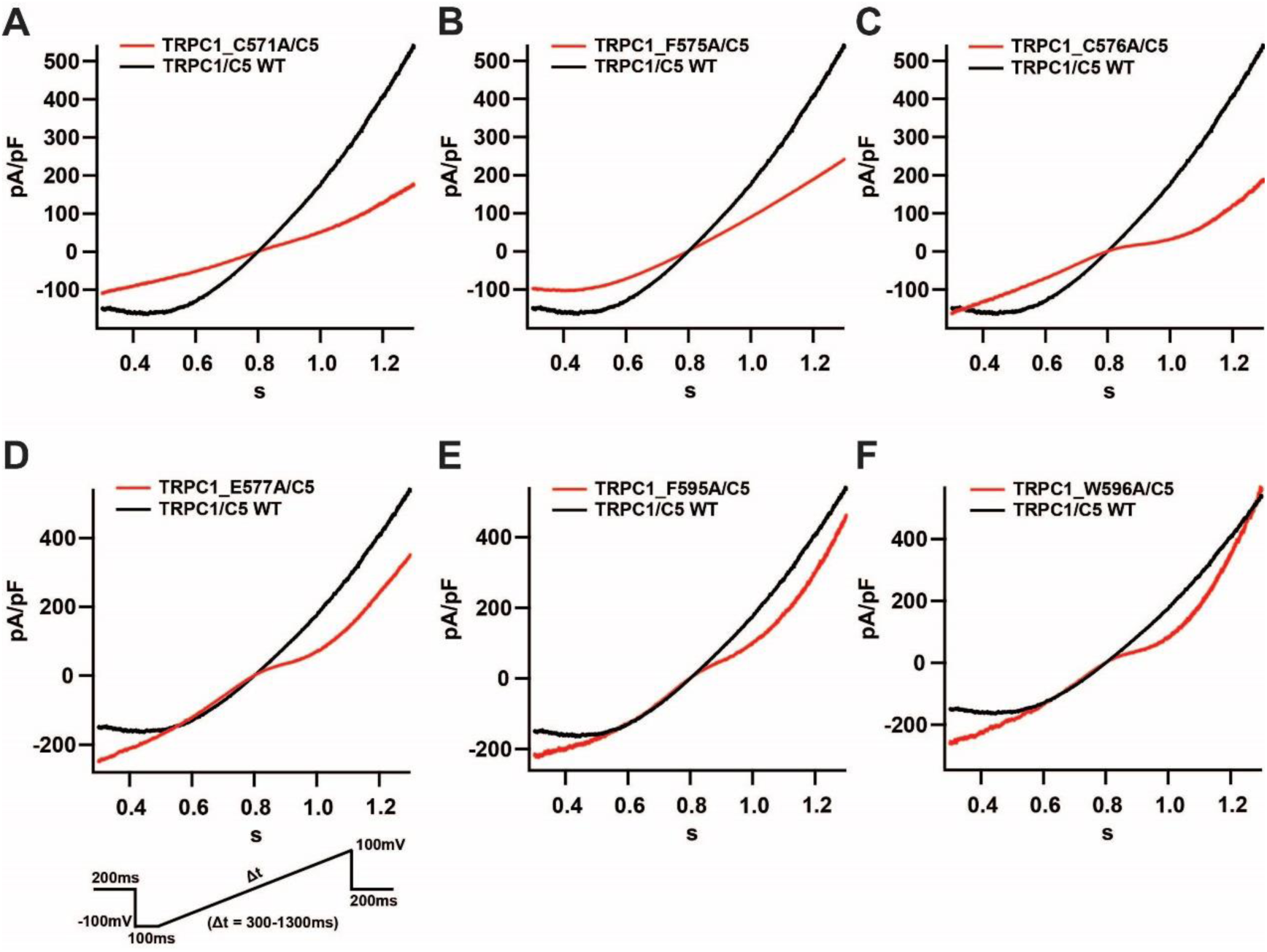
The influence of TRPC1’s pore-loop and poor-helix mutations on TRPC1/5 heteromer activation. **(A to F)** The characteristic current curves of TRPC1/5 heteromer caused by various mutations in the TRPC1 subunit (red) compared to TRPC1/5 wild type (WT) (black).

**Figure S7.**
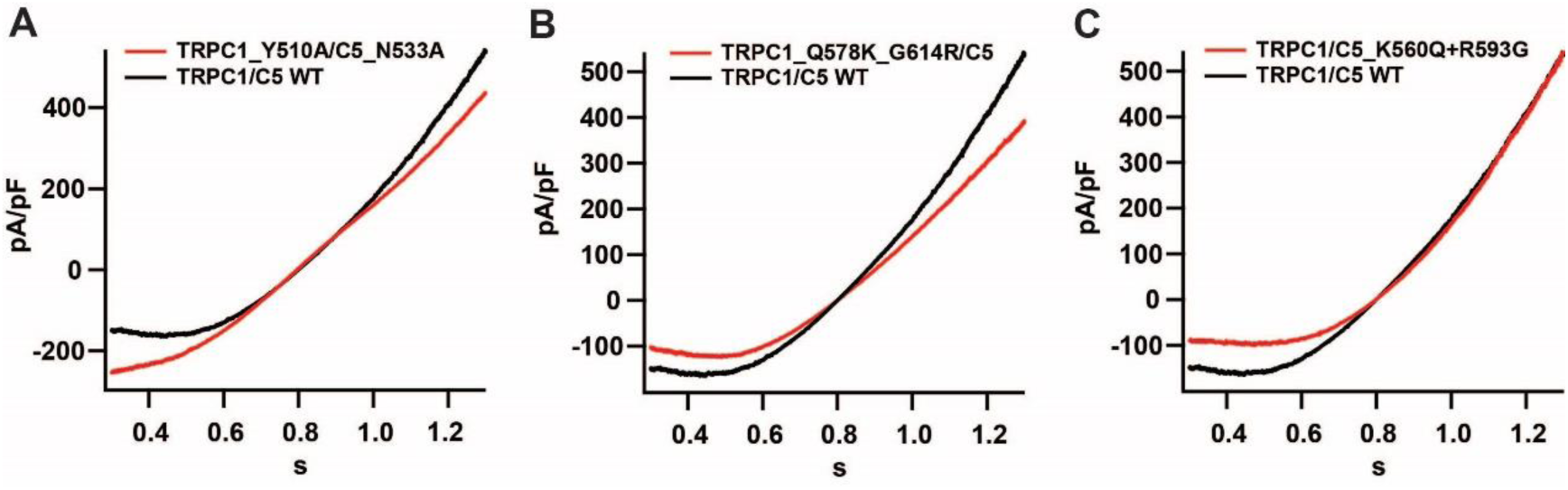
The influence of TRPC1-TRPC5 interface mutations on TRPC1/5 heteromer activation. **(A to C)** The characteristic current curves of TRPC1/5 heteromer caused by various mutations in the TRPC1 or TRPC5 subunit (red) compared to TRPC1/5 wild type (WT) (black).

**Figure S8.**
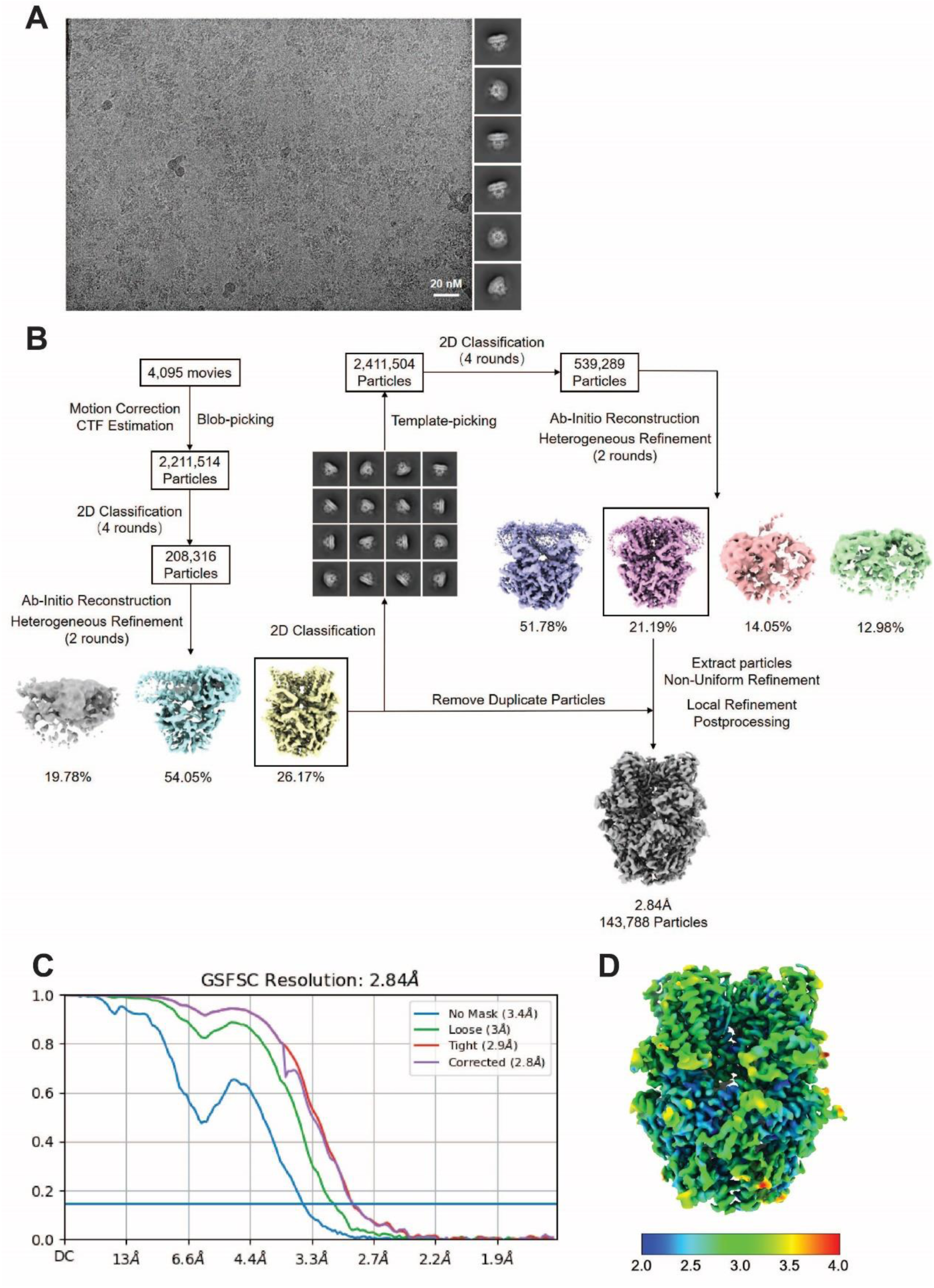
Flow chart for TRPC1/5 cryo-EM data processing. **(A)** Representative of micrograph from 4,095 movies of TRPC1/5 sample in vitreous ice (left) and 2D class averages (right). Scale bar is at 20 nm. **(B)** Flow chart of the cryo-EM data processing procedure. Representative image of 3D reconstructions from final non-uniform refinement and local refinement, and the final density map of TRPC1/5 heteromer with overall resolution of 2.84 Å. **(C)** Gold-standard Fourier shell correlation (GSFSC) curves for the 3D reconstruction of TRPC1/5 heteromer. **(D)** Local resolution estimation.

**Table S1.** Cryo-EM data collection, refinement and validation statistics.

|  |  |
| --- | --- |
| Structure | TRPC1/5 |
| EMDB accession code | (EMD-62060) |
| PDB accession code | (PDB 9K4I) |
| **Data collection and processing** |  |
| Magnification | 105,000 |
| Voltage (kV) | 300 |
| Electron exposure (e-/Å<sup>2</sup>) | 54 |
| Defocus range (μm) | -1.0 ~ -2.0 |
| Pixel size (Å) | 0.83 |
| Symmetry imposed | *C1* |
| Initial particle images (#) | 2,411,504 |
| Final particle images (#) | 143,788 |
| Map resolution (Å) | 2.84 |
| FSC threshold | 0.143 |
| **Refinement** |  |
| Initial model used (PDB code) | 6AEI |
| Model resolution (Å) | 2.89 |
| FSC threshold | 0.143 |
| **Model composition** |  |
| Non-hydrogen atoms | 22,192 |
| Protein residues | 2709 |
| Ligands | 10 |
| **B factors (Å<sup>2</sup>)** |  |
| Protein | 72.71 |
| Ligand | 76.54 |
| r.m.s. deviations |  |
| Bond lengths (Å) | 0.008 |
| Bond angles (°) | 0.944 |
| **Validation** |  |
| MolProbity score | 2.01 |
| Clashscore | 17 |
| Poor rotamers (%) | 0 |
| Ramachandran plot |  |
| Favored (%) | 94.46 |
| Allowed (%) | 5.54 |
| Disallowed (%) | 0 |

## References

1. D. E. Clapham, TRP channels as cellular sensors. Nature 426, 517–524 (2003)

2. Venkatachalam K, Montell C. TRP channels. Annu Rev Biochem. 76, 387–417 (2007)

3. Montell C, Birnbaumer L, Flockerzi V, The TRP channels, a remarkably functional family. Cell 108, 595–598 (2002)

4. Freichel M, Vennekens R, Olausson J, Stolz S, Philipp SE, Weissgerber P, Flockerzi V, Functional role of TRPC proteins in native systems: implications from knockout and knock-down studies. J Physiol. 567, 59–66 (2005)

5. Fowler MA, Sidiropoulou K, Ozkan ED, Phillips CW, Cooper DC, Corticolimbic expression of TRPC4 and TRPC5 channels in the rodent brain. PLoS ONE 2, e573 (2007)

6. Strübing C, Krapivinsky G, Krapivinsky L, Clapham DE, TRPC1 and TRPC5 form a novel cation channel in mammalian brain. Neuron 29, 645–655 (2001)

7. Strübing C, Krapivinsky G, Krapivinsky L, Clapham DE, Formation of novel TRPC channels by complex subunit interactions in embryonic brain. J. Biol. Chem. 278, 39014–39019 (2003)

8. Kollewe A, Schwarz Y, Oleinikov K, Raza A, Haupt A, Wartenberg P, Wyatt A, Boehm U, Ectors F, Bildl W, Zolles G, Schulte U, Bruns D, Flockerzi V, Fakler B, Subunit composition, molecular environment, and activation of native TRPC channels encoded by their interactomes. Neuron 110, 4162–4175 (2022)

9. Greka A, Navarro B, Oancea E, Duggan A, Clapham DE, TRPC5 is a regulator of hippocampal neurite length and growth cone morphology. Nat. Neurosci. 6, 837–845 (2003)

10. Hartmann J, Dragicevic E, Adelsberger H, Henning HA, Sumser M, Abramowitz J, Blum R, Dietrich A, Freichel M, Flockerzi V, Birnbaumer L, Konnerth A TRPC3 channels are required for synaptic transmission and motor coordination. Neuron 59, 392–398 (2008)

11. Riccio A, Li Y, Moon J, Kim KS, Smith KS, Rudolph U, Gapon S, Yao GL, Tsvetkov E, Rodig SJ, Van’t Veer A, Meloni EG, Carlezon WA Jr, Bolshakov VY, Clapham DE, Essential role for TRPC5 in amygdala function and fear-related behavior. Cell 137: 761–772 (2009)

12. Riccio A, Li Y, Tsvetkov E, Gapon S, Yao GL, Smith KS, Engin E, Rudolph U, Bolshakov VY, Clapham DE, Decreased anxiety-like behavior and Galphaq/11-dependent responses in the amygdala of mice lacking TRPC4 channels. J. Neurosci. 34, 3653–3667 (2014)

13. Cooper, D, Klipec, W, Deeney, B, Williamson, C, & Ostertag, E, Deletion of the rat trpc4 gene and its influence on motivated responding for natural reward. Nature Precedings, 41, 138–155 (2012)

14. Phelan KD, Mock MM, Kretz O, Shwe UT, Kozhemyakin M, Greenfield LJ, Dietrich A, Birnbaumer L, Freichel M, Flockerzi V, Zheng F, Heteromeric canonical transient receptor potential 1 and 4 channels play a critical role in epileptiform burst firing and seizure-induced neurodegeneration. Mol. Pharmacol. 81, 384–392 (2012)

15. Phelan KD, Shwe UT, Abramowitz J, Wu H, Rhee SW, Howell MD, Gottschall PE, Freichel M, Flockerzi V, Birnbaumer L, Zheng F, Canonical transient receptor channel 5 (TRPC5) and TRPC1/4 contribute to seizure and excitotoxicity by distinct cellular mechanisms. Mol. Pharmacol. 83, 429–438 (2013)

16. Chu WG, Wang FD, Sun ZC, Ma SB, Wang X, Han WJ, Wang F, Bai ZT, Wu SX, Freichel M, Xie RG, Luo C, TRPC1/4/5 channels contribute to morphine-induced analgesic tolerance and hyperalgesia by enhancing spinal synaptic potentiation and structural plasticity. FASEB J. 34, 8526–8543 (2020)

17. Gupta V, Ben-Mahmoud A, Ku B, Velayutham D, Jan Z, Yousef Aden A, Kubbar A, Alshaban F, Stanton LW, Jithesh PV, Layman LC, Kim HG, Identification of two novel autism genes, *TRPC4* and *SCFD2*, in Qatar simplex families through exome sequencing. Front Psychiatry. 14, 1251884 (2023)

18. Li Y, Cacciottolo TM, Yin N, He Y, Liu H, Liu H, Yang Y, Henning E, Keogh JM, Lawler K, Mendes de Oliveira E, Gardner EJ, Kentistou KA, Laouris P, Bounds R, Ong KK, Perry JRB, Barroso I, Tu L, Bean JC, Yu M, Conde KM, Wang M, Ginnard O, Fang X, Tong L, Han J, Darwich T, Williams KW, Yang Y, Wang C, Joss S, Firth HV, Xu Y, Farooqi IS, Loss of transient receptor potential channel 5 causes obesity and postpartum depression. Cell 187, 4176–4192 (2024)

19. Storch U, Forst AL, Philipp M, Gudermann T, Mederos y Schnitzler M, Transient receptor potential channel 1 (TRPC1) reduces calcium permeability in heteromeric channel complexes. J. Biol. Chem. 287, 3530–3540 (2012)

20. Kim J, Ko J, Myeong J, Kwak M, Hong C, So I, TRPC1 as a negative regulator for TRPC4 and TRPC5 channels. Pflugers Arch. 471, 1045–1053 (2019)

21. Duan J, Li J, Chen GL, Ge Y, Liu J, Xie K, Peng X, Zhou W, Zhong J, Zhang Y, Xu J, Xue C, Liang B, Zhu L, Liu W, Zhang C, Tian XL, Wang J, Clapham DE, Zeng B, Li Z, Zhang J, Cryo-EM structure of TRPC5 at 2.8-Å resolution reveals unique and conserved structural elements essential for channel function. Sci. Adv. 5, eaaw7935 (2019)

22. Vinayagam D, Mager T, Apelbaum A, Bothe A, Merino F, Hofnagel O, Gatsogiannis C, Raunser S, Electron cryo-microscopy structure of the canonical TRPC4 ion channel. Elife. 7, e36615 (2018)

23. Duan J, Li J, Zeng B, Chen GL, Peng X, Zhang Y, Wang J, Clapham DE, Li Z, Zhang J, Structure of the mouse TRPC4 ion channel. Nat. Commun. 9, 3102 (2018)

24. Song K, Wei M, Guo W, Quan L, Kang Y, Wu JX, Chen L, Structural basis for human TRPC5 channel inhibition by two distinct inhibitors. Elife. 10, e63429 (2021)

25. Yang Y, Wei M, Chen L. Structural identification of riluzole-binding site on human TRPC5. Cell Discov. 8, 67 (2022)

26. Jumper J, Evans R, Pritzel A, Green T, Figurnov M, Ronneberger O, Tunyasuvunakool K, Bates R, Žídek A, Potapenko A, Bridgland A, Meyer C, Kohl SAA, Ballard AJ, Cowie A, Romera-Paredes B, Nikolov S, Jain R, Adler J, Back T, Petersen S, Reiman D, Clancy E, Zielinski M, Steinegger M, Pacholska M, Berghammer T, Bodenstein S, Silver D, Vinyals O, Senior AW, Kavukcuoglu K, Kohli P, Hassabis D, Highly accurate protein structure prediction with AlphaFold. Nature 596, 583–589 (2021)

27. Varadi M, Bertoni D, Magana P, Paramval U, Pidruchna I, Radhakrishnan M, Tsenkov M, Nair S, Mirdita M, Yeo J, Kovalevskiy O, Tunyasuvunakool K, Laydon A, Žídek A, Tomlinson H, Hariharan D, Abrahamson J, Green T, Jumper J, Birney E, Steinegger M, Hassabis D, Velankar S, AlphaFold Protein Structure Database in 2024: providing structure coverage for over 214 million protein sequences, Nucleic Acids Res. 52, D368–D375 (2024)

28. Pettersen EF, Goddard TD, Huang CC, Couch GS, Greenblatt DM, Meng EC, Ferrin TE, UCSF Chimera--a visualization system for exploratory research and analysis. J. Comput. Chem. 25, 1605–12 (2004)

29. Emsley P, Lohkamp B, Scott WG, Cowtan K. Features and development of Coot. Acta Crystallogr D Biol Crystallogr. 66, 486–501 (2010)

30. Afonine PV, Poon BK, Read RJ, Sobolev OV, Terwilliger TC, Urzhumtsev A, Adams PD. Real-space refinement in PHENIX for cryo-EM and crystallography, Acta Crystallogr D Struct Biol. 74, 531–544 (2018)

